# Syntactic Constructions Drive Cortical Tracking in the Absence of Lexical Content: An Electrophysiological Investigation of Sentence Processing During Reading

**DOI:** 10.1101/2023.07.17.549412

**Authors:** M. Blake Rafferty, Tim Saltuklaroglu, Eun Jin Paek, Kevin Reilly, David Jenson, David Thornton, Devin M. Casenhiser

**Affiliations:** New Mexico State University, Las Cruces, NM; University of Tennessee Health Science Center, Knoxville, TN; Washington State University, Spokane, WA; University of Oslo, Norway

**Keywords:** neural synchrony, oscillations, EEG, syntax, language comprehension

## Abstract

It has been suggested that the synchronization of neural oscillations to syntactic units, such as phrases or clauses, is dependent on lexically-derived projections of syntactic structure. This assertion is based on recent evidence that participants are unable to effectively track syntax when listening to jabberwocky sentences, in which content words are replaced with pseudowords thereby eliminating lexically-derived syntactic projections (Coopmans et al., 2022; Kaufeld et al., 2020). In the present study, we extend the findings from these two studies and present evidence that participants can in fact track syntactic units in jabberwocky sentences when the stimuli are presented visually – a methodological difference that allows participants to easily parse individual words in the sentence. We interpret this finding as indicating that tracking the phrase structure of a sentence can take place in the absence of content words and does not crucially depend on their lexical projections.

## Introduction

It has been suggested that the firing of neural oscillations can temporally synchronize with the occurrence of multi-unit linguistic representations (e.g., syntactic constituents such as phrases or clauses) (Martin, 2020; Martin & Doumas, 2017; Meyer et al., 2019; Meyer et al., 2020). Experimental evidence for this was first demonstrated by Ding and colleagues (Ding, Melloni, Tian, et al., 2017; Ding, Melloni, Yang, et al., 2017; Ding et al., 2016) using a frequency-tagging paradigm in which participants’ neural responses to monosyllabic words were enhanced when word boundaries co-occurred with phrase and sentence boundaries.

In subsequent work, Kaufeld et al. (2020) and Coopmans et al. (2022) further explored the effect in naturalistic speech by measuring the amount mutual information (MI) found between the EEG signal and syntactic annotations denoting phrase boundaries. Both studies demonstrated that the effect could be found in forward audio sentences, but not in sentences in which the audio had been reversed. Coopmans et al. additionally demonstrated that while sentences show the highest level of neural synchrony at phrase boundaries, the difference was not significantly greater than for idioms and syntactic prose (syntactically correct, but semantically anomalous sentences). Conversely, sentences evidenced significantly higher neural synchrony to phrase boundaries than did word lists or jabberwocky (sentences wherein content words are replaced with pseudowords while function words and morphemes are retained). Jabberwocky sentences, in fact, did not differ from word lists in the amount of neural synchrony measured at phrase boundaries. This last finding is of particular interest to the current study since it suggests that parsing of sentence structure may be hindered due to the lack of content words. Coopmans et al. interpret their results as being consistent with a lexicalist framework in which a sentence’s argument structure is projected from lexical items: “Jabberwocky sentences are structured sequences that contain both function words and inflectional morphology, but they lack content words and therefore miss the information carried by their argument structure…The lexical-syntactic difference between regular and jabberwocky sentences thus explains why they elicit different degrees of phrase-level speech tracking” (Coopmans et al., 2023, p. 406).

The approach to syntax wherein argument structure is thought to project from lexical items to syntax is broadly referred to as lexicalism. The notion can be traced back to Chomsky’s (Chomsky et al., 1970) Lexicalist Hypothesis, although several variations of the hypothesis (e.g., Aronoff, 1976; Bresnan, 1980; Di Sciullo & Williams, 1987; Lapointe, 1980) as well as both theoretical and psycholinguistic models of language comprehension have been proposed since Chomsky’s initial formulation (e.g., Culicover & Jackendoff, 2005; Hagoort, 2013; Kaplan & Bresnan 1981; MacDonald et al., 1994; Matchin & Hickock 2019; Pollard & Sag, 1994). According to the standard lexicalist approach, lexical processes generate output that serves as the input to syntactic processes. These processes are separate and unidirectional (i.e., syntactic processes do not feed lexical ones).

Despite its popularity, the lexicalist approach is not universally accepted among language researchers. Adjudication of the specific criticisms of the lexicalist approach is beyond the scope of this paper; instead, we summarize by saying that the arguments against the approach have disputed the basic facts upon which the lexicalist model is predicated and/or appealed to parsimony by arguing that separate lexical and syntactic processes are not required to account for the data (for details see, e.g., Bruening, 2018; Goldberg, 1995; Halle & Marantz 1993; Haspelmath, 2011; Jackendoff, 2017; Krauska & Lau 2023).

If Coopman’s et al.’s interpretation of their results is accurate, their study would provide useful psycholinguistic support in favor of a lexicalist hypothesis. However, there is some recent evidence produced in our lab (Rafferty et al., 2023) that appears to be at odds with the interpretation of the results of Coopmans et al. In this study, we used a minimal phrase paradigm to investigate whether individual noun phrases composed of a determiner and a pseudoword were sufficient to elicit neural synchrony given the lack of lexical semantics. This study recorded EEG activity while participants read mixed blocks of syntactically well-formed phrases composed of a determiner and a pseudoword (*the moop*) and a wordlist condition composed of the same pseudoword followed by a determiner (*moop the*). Results showed that in comparison to the words condition, there was significantly greater neural synchrony for the phrase condition (measured by inter-trial phase coherence) at the phrase rate with no difference between conditions at the word rate. This finding suggests that the minimal phrases were sufficient to induce neural synchrony even when they contained no content words. Since the pseudowords do not contain lexical projections, it would seem difficult to reconcile the results of this study with those of Kaufeld et al. and Coopmans et al. The findings furthermore appear to provide evidence counter to a lexicalist interpretation.

Accordingly, we suggest an alternative interpretation of the results obtained by Kaufeld et al. and Coopmans et al. Namely, that their participants crucially had difficulty tracking utterances at the word level resulting in their apparent lack of phrase-level tracking. In fact, Kaufeld et al. report that tracking at the word rate is significantly better for sentences than for jabberwocky (β = 0.48; *p* < 0.01) and nearly significantly worse for jabberwocky than for word lists (β = −0.33; *p* = 0.06). In comparison, tracking at the word rate between sentences and the word list condition shows no significant difference (β = 0.16; *p* = 0.14). Thus, the reduced phrase tracking observed in jabberwocky may be accounted for by participants’ difficulties parsing individual words from the speech stream. That is, if individuals have difficulty parsing individual words from the speech stream, they would not be able to accurately track phrases.

To further clarify whether tracking of phrases requires lexically derived projections to syntax, we extend the findings of Kaufeld et al. and Coopmans et al. by investigating whether syntactic frames would elicit synchrony in the absence of lexical content words. To remove the difficulties associated with parsing individual words from the auditory stream, we opted to use visually presented stimuli. Sentence stimuli were composed of syntactically identical English and jabberwocky sentences using the English caused-motion construction (Goldberg, 1995). We chose the caused motion construction because of the high cue validity of its syntactic structure and meaning (cf. Goldberg 2006; Goldberg et al. 2005). Its pattern is represented below using simplified constructionist conventions. Other representational conventions may work as well, and there is nothing on the phrase level to differentiate this account from a standard generativist approach since phrasal boundaries would occur in the same location in either approach. This is because complex constructions may be composed of smaller constructions. In this case, the caused motion construction is composed of Subject-Predicate, NP, VP, and PP constructions.

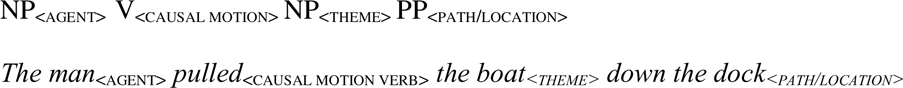

To control for effects related to semantic composition in the English sentences, participants read scrambled wordlists with the same lexical items, constructed to minimize coherent phrase structure. To control for both semantics and syntax at once, we also included a pseudoword wordlist control condition. All stimuli were presented visually using rapid serial visual presentation to ensure that between-condition differences in phrase-level tracking did not result from ambiguous word boundaries in the items containing unfamiliar pseudowords. Examples of the stimuli for each condition are listed in *Table 1*.

**Table 1:** Example stimuli from each condition.

|  | **Example Stimuli** |
| --- | --- |
| **English Sentence** | the man pulled the boat down the dock |
| **Jabberwocky Sentence** | the dex mooped the pob down the jarm |
| **English Wordlist** | boat the the man dock down pulled the |
| **Pseudoword Wordlist** | pob the the dex jarm down mooped the |

**Table 2:** Estimated marginal means from analysis of log-transformed MI values for tracking syntactic node closure.

| **Contrasts** | **Estimate** | **SE** | **df** | ***t*-values** | ***p*-values** |
| --- | --- | --- | --- | --- | --- |
| English Sentence – English Wordlist | 0.61 | 0.11 | 48 | 5.39 | **<0.001** |
| English Sentence – Jabberwocky Sentence | -0.06 | 0.13 | 45.2 | -0.43 | 0.69 |
| English Sentence – Pseudoword Wordlist | 0.75 | 0.13 | 24 | 5.85 | **<0.01** |
| English Wordlist – Jabberwocky Sentence | -0.66 | 0.13 | 24 | -5.05 | **<0.01** |
| English Wordlist – Pseudoword Wordlist | 0.14 | 0.13 | 45.2 | 1.11 | 0.27 |
| Jabberwocky Sentence – Pseudoword Wordlist | 0.80 | 0.11 | 48 | 7.16 | **<0.01** |

**Table 3:** Estimated marginal means from analysis of log-transformed MI values for tracking syntactic node count.

| **Contrasts** | **Estimate** | **SE** | **df** | ***t*-values** | ***p*-values** |
| --- | --- | --- | --- | --- | --- |
| English Sentence – English Wordlist | 0.71 | 0.12 | 48 | 6.09 | **<0.01** |
| English Sentence – Jabberwocky Sentence | -0.06 | 0.13 | 54 | -0.44 | 0.66 |
| English Sentence – Pseudoword Wordlist | 0.95 | 0.13 | 54 | 7.25 | **<0.01** |
| English Wordlist – Jabberwocky Sentence | -0.77 | 0.13 | 54 | -5.85 | **<0.01** |
| English Wordlist – Pseudoword Wordlist | 0.24 | 0.13 | 54 | 1.84 | 0.08 |
| Jabberwocky Sentence – Pseudoword Wordlist | 1.01 | 0.12 | 48 | 8.65 | **<0.01** |

**Table 4:** Estimated marginal means from analysis of EEG power for tracking words.

| **Contrasts** | **Estimate** | **SE** | **df** | ***t*-values** | ***p*-values** |
| --- | --- | --- | --- | --- | --- |
| English Sentence – English Wordlist | 0.02 | 0.01 | 39 | 2.41 | **0.02** |
| English Sentence – Jabberwocky Sentence | 0.04 | 0.01 | 44.9 | 4.28 | **<0.01** |
| English Sentence – Pseudoword Wordlist | 0.06 | 0.01 | 24 | 4.96 | **<0.01** |
| English Wordlist – Jabberwocky Sentence | 0.01 | 0.01 | 24 | 1.17 | 0.25 |
| English Wordlist – Pseudoword Wordlist | 0.03 | 0.01 | 44.9 | 4.00 | **<0.01** |
| Jabberwocky Sentence – Pseudoword Wordlist | 0.02 | 0.01 | 39 | 2.18 | **0.04** |

As in Kaufeld et al. and Coopmans et al., we quantified the degree of neural synchrony as the degree of MI between EEG recordings and the syntactic structure of the sentences. Should the composition of syntactic structure necessitate the presence of lexical-syntactic projections, English sentences would elicit greater synchrony (as evidenced by greater MI) than all other conditions. We hypothesize, however, that phrase-level tracking can take place even when sentences lack content words. As a result, we anticipate that both English and jabberwocky sentences should elicit greater MI than both English and pseudoword wordlist conditions.

To preview our results, we find that participants were able to track phrases in both English and jabberwocky sentences to a similar degree, whereas considerably less phrase-level tracking was observed for the two wordlist conditions. These results are consistent with prior studies showing that neural oscillations may become synchronized with the boundaries of syntactic phrases. However, unlike previous work, our findings suggest that phrase boundaries may be inferred even in the absence of lexical projections from content words.

## Method

### Participants

25 native English speakers (16 females & 9 males) aged 20-34 (M = 24.32; SD = 3.69) were recruited for this study. Participants had normal or corrected to normal vision and no reported history of neurological injury or pathology. All participants were right-handed as assessed by the Edinburgh Handedness Inventory (Oldfield, 1971). This study was approved by the Institutional Review Board of the University of Tennessee Health Science Center (Review Number: 17-04047-XP). Prior to the experiment, participants provided signed informed consent on a document approved by the Institutional Review Board.

### Materials

Four sets of sentence stimuli were constructed. The first set consisted of 20 grammatical English sentences that adhered to the structure of the caused-motion construction (e.g., *The man pulled the boat down the dock).* That is, each sentence included a subject noun phrase, and a verb phrase with direct object and prepositional phrase. Subject NPs contained exclusively animate nouns (e.g., *child*, *snail, team*) to facilitate the assignment of the thematic role _<AGENT>_. For VPs, all verbs were action verbs (e.g., *tossed*, *pushed*, *flipped*) to encourage motion-based interpretations (i.e., to facilitate the assignment of a _<CAUSAL MOTION>_ argument to the verb). All verbs were also morphologically regular and remained single syllable when marked for past tense. To satisfy the _<PATH/LOCATION>_ argument, PPs consisted of one of five prepositions (*down, through*, *off*, *up*, or *to*) and an NP containing an inanimate noun which, when paired with the preposition, may be interpreted as a location (e.g., *through the air, down the road, up the trail*). All NPs followed the structure *determiner + noun*, regardless of the position of the NP in the sentence, and the indefinite article *the* was used as the sole determiner. The full list of English sentence stimuli is included within the Supplementary Materials.

In addition to the English sentence condition, we also constructed 20 lexically matched English wordlists by pseudo-randomizing order of the words within the English sentences to reduce global syntactic structure. All words were a single syllable, morphologically regular, and no longer than 7 letters. We then constructed 20 jabberwocky sentences, which contained the same syntactic structure as the English caused-motion sentences but wherein content words were replaced with pseudowords. To control for lexical-semantics and syntactic structure, we also constructed a pseudoword wordlist condition. Pseudowords were created by swapping one letter from real English words. We controlled for phonotactic probability by first calculating the summed phonetic probability and bi-phone probability for each pseudoword using the online phonotactic probability calculator from the University of Kansas (Vitevitch & Luce, 2004). Then we compared these summed totals (M = 1.173, SD = 0.0605) to their real-English counterparts (M = 1.176, SD = 0.0508) using paired-samples t-tests. No significant effects were found, *t*(1,59) = −0.5158, p = 0.608. Importantly, the lack of statistical effect does not entirely rule out the possibility that phonotactic differences may contribute to the findings reported here (Sassenhagen & Alday, 2016). Nonetheless, these results illustrate that the phonotactic probabilities for pseudowords largely overlapped with those for the real words included in this study.

### Procedure

For EEG recordings participants sat in a comfortable chair approximately 50 inches away from a television monitor in a magnetically shielded, double-walled, sound-attenuating booth. Stimuli were presented visually to reduce potential difficulties associated with parsing unfamiliar pseudowords from continuous speech.

Each trial was preceded by a 1000ms period during which the monitor screen showed a fixation cross in its center to bring participants’ focus to the center of the screen and eliminate any saccadic activity associated with trial onset. After this period, the fixation cross disappeared, and sentences were presented one word at a time. Each word was visible on the screen for 500ms, after which it was immediately replaced by the next word. Participants were instructed to read the sentences silently. 1000ms after the offset of the sentence-final word, participants heard a beep to indicate that they should press a button on a hand-held controller to advance to the next trial. For an example timeline, see *Figure 1*.

**Figure 1:**
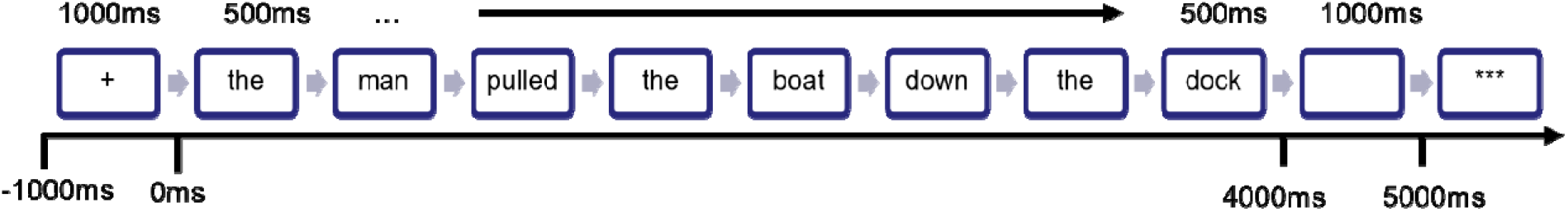
Trial timeline. The top row denotes the duration of each item, and the bottom row indicates when each item appears relative to sentence onset. At the beginning of each trial, a fixation cross appeared for 1000ms. After this, words appeared on the monitor one at a time for 500ms each. Following the final word of each trial, there was a 1000ms blank screen followed by the appearance of three stars that indicated participants should press a button on a handheld keypad to initiate the next trial.

The experimental procedure was divided into eight mixed blocks. Each block consisted of 40 randomized experimental trials, with 10 from each condition. No more than three items from one condition occurred in sequence, and block order was randomized across participants. Blocks lasted approximately 7 minutes each, for a total of approximately 1 hour in the EEG booth.

### EEG Acquisition

EEG data were recorded at a sampling rate of 1000Hz using 64 electrodes mounted on an unlinked and sintered NeuroScan QuikCap 64, configured according to the extended international 10-20 system (Klem et al., 1999). Horizontal eye movement was recorded by placing electrodes on the lateral canthi of each eye (horizontal electrooculography—HEOG); meanwhile, vertical eye movement was recorded by placing electrodes on the superior and inferior orbit (vertical electrooculography VEOG) of the left eye. Electrocardiography (EKG) was recorded from bipolar electrodes placed on both sides of participants’ neck once their pulse was located.

### EEG Pre-processing

Data were preprocessed offline using the open-source FieldTrip toolbox (v20200801; Oostenveld et al., 2011) in MATLAB (The MathWorks, Inc., Matick, MA, USA). Continuous EEG data were first bandpass filtered between 0.1 and 100 Hz (1st order, two-pass Butterworth), and 60Hz line-noise was then removed using the fast-Fourier transform. Data were then epoched from −3000ms to +6000ms relative to stimulus onset. Trials with response times that were less than 100ms were removed. Next, we visually inspected the single trial data and removed noisy channels (M = 1.6; SD = 1.32). Data were then re-referenced to the common average, and trials with clear movement artifacts were removed. We removed ocular and cardiac artifacts using independent components analysis. We then applied the resultant component weights to the original data and subtracted the artifacts. On average 2.6 components were removed per participant. Using the cleaned data, we interpolated noisy channels previously removed using the ‘spline’ method. We conducted a final manual inspection of all trials and removed any remaining artifactual trials. On average, subjects maintained > 58 trials per condition (SD = 10.95).

### Mutual Information

#### Tracking Node Closure

To investigate whether predictable syntactic frames would elicit neural synchrony, we calculated mutual information (MI) between the EEG and vector annotations of abstract syntactic structure at the frequency of occurrence for phrases (.75Hz) (Brodbeck et al., 2018; Coopmans et al., 2022; Kaufeld et al., 2020). We selected MI for comparability to Kaufeld et al. and Coopmans et al. Pairing MI with annotations of sentence structure allowed us to assess cortical tracking of phrases that occur non-metrically. Annotations were created manually by first identifying all time samples that corresponded to phrase final words in each sentence. We then coded each of these samples as ones to represent the closure of syntactic constituents or “nodes”. All other samples were coded as zeroes (see *Figure 2A*). The same annotations of syntactic structure were evaluated in the MI analysis for all conditions under the assumption that cortical responses to wordlists should demonstrate low MI with the structure annotations since the wordlists do not contain internal syntactic structure.

**Figure 2:**
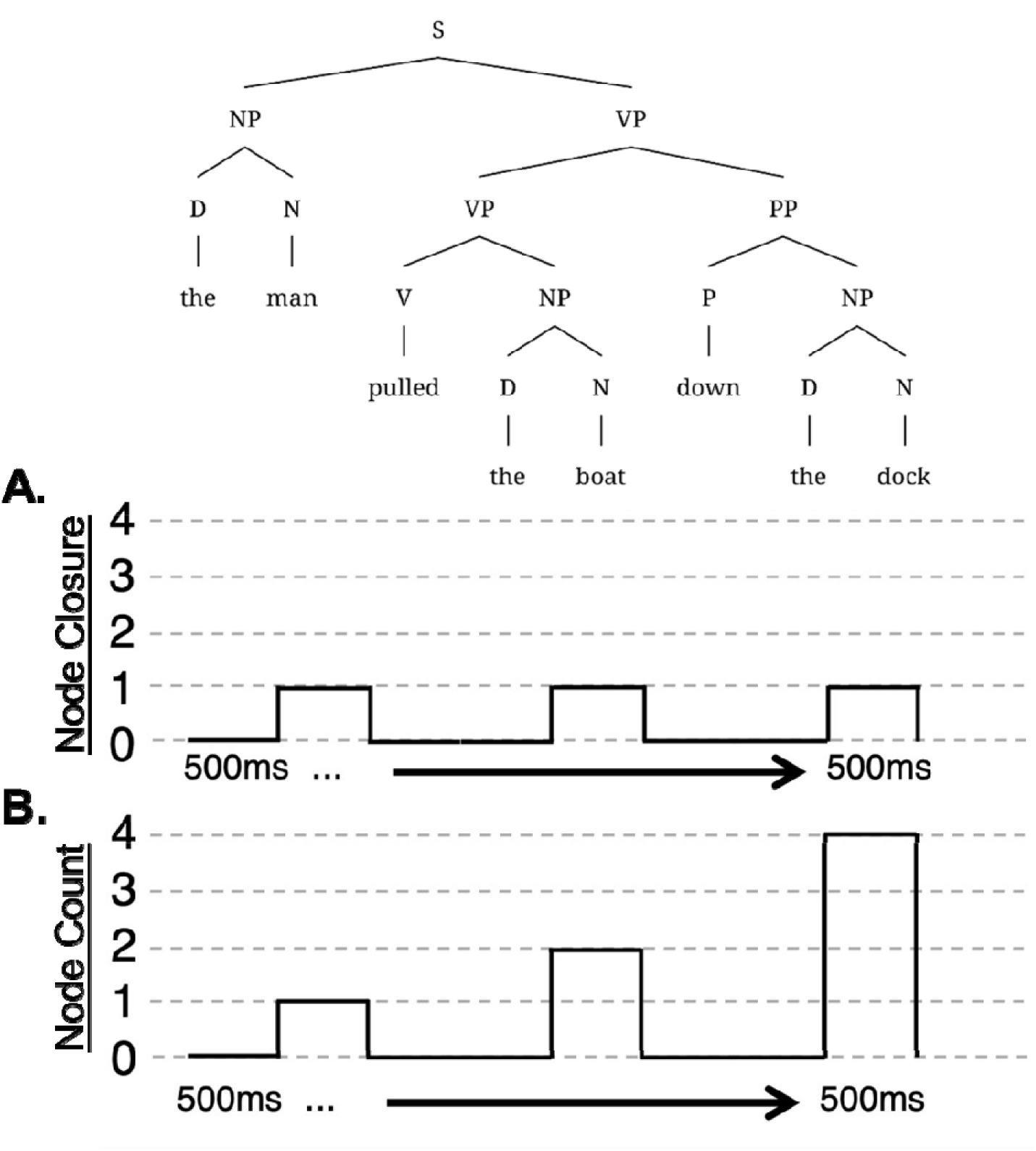
A. Visual depictions of annotations that encode the closure of syntactic nodes according to the phrase structure of the sentence. B. Annotations that encode the number of syntactic phrasal nodes that are closed by a given item (i.e., “node count”) according to the phrase structure of the sentence.

To ensure that unequal trial counts did not bias mutual information values, we randomly selected an equal number of trials across all conditions based upon the minimum number of trials remaining for each condition. Following this, we band-pass filtered the data into frequencies corresponding to the rate of occurrence for phrases (0.6 - 0.9Hz, centered around the phrase rate of .75Hz), separately for each condition (3^rd^ order, two-pass Butterworth). We then Hilbert-transformed the data and normalized the resulting real and imaginary-valued coefficients separately using the Gaussian copula method (Ince et al., 2017). The resulting phase and power time series for each trial were then recombined and trimmed to be equal in length to the syntactic annotations described above, which were added to the data structures for each condition as “dummy” sensors. We then concatenated all trials end-to-end and calculate MI for each condition as:

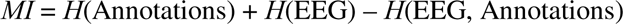

where *H*(Annotations) is the Shannon entropy of the Hilbert transformed annotations of syntactic structure, *H*(EEG) is the entropy of the Hilbert transformed EEG, and *H*(EEG, Annotations) is their joint entropy.

Prior studies using speech stimuli have calculated MI within fixed sets of electrodes and at speech-brain lags that correspond to the processing window for auditory responses (e.g., Coopmans et al., 2022; Gross et al., 2013; Kaufeld et al., 2020; Keitel et al., 2017; Keitel et al., 2018). However, the topography and time-course of cortical tracking responses to visually presented stimuli as implemented in the current study is unknown. To ensure the selection of appropriate time windows and electrodes for analysis, we calculated the temporal mutual information function for each condition (TMIF; De Clercq et al., 2023). This consisted of convolving the EEG data with the structural annotations one sample at a time over speech-brain lags ranging −300ms – 500ms, relative to stimulus onset. Following this, we concatenated the resulting values into a time-series (i.e., the TMIF) to assess the time-course of cortical responses between conditions. We then log-transformed the data and removed outlier participants whose average responses were in the 2.5^th^ and 97.5^th^ percentiles.

#### Tracking Node Count

While our analysis of node closure captures the formation of syntactic phrases at the local level, syntactic structure encodes information about hierarchical relationships amongst multiple constituents across the length of a sentence (Brennan & Hale, 2019). Thus, in addition to assessing cortical tracking of node closures, we also computed MI between the EEG and annotations that encode the accumulated number of syntactic constituents that each phrase-final word completes—i.e., node count (Brennan & Hale, 2019; Coopmans et al., 2022; Lo et al., 2022; Nelson et al., 2017). For this, we copied the annotations created above and coded all non-zero samples with the number of phrasal nodes that may be closed at that point in time. Visual depictions of the annotations corresponding to node count can be seen in *Figure 2B*.

With this analysis, we investigated the possibility that the presence of intact grammatical structure in jabberwocky items may allow for phrase formation at the local level, but utterance-spanning constituents may require the ability to associate thematic roles with specific items. If this is the case, we expected that MI for English and jabberwocky sentences would not differ when considering node closure but would differ when considering node count. Alternatively, the semantics associated with the caused-motion construction may license the formation of such hierarchical syntactic relationships. If this is the case, we anticipated no differences between English and jabberwocky sentences related to tracking node count.

#### Statistical Testing

To select appropriate temporal and topographic parameters for our analysis of cortical tracking, we performed one-tailed omnibus analyses of variance using cluster-based permutation tests on the TMIFs for node closure and node count, separately (Maris & Oostenveld, 2007). This procedure involved computing an *F*-statistic between conditions for all samples and electrodes. Sample-electrode pairs that exceed a pre-specified threshold (*p <* .05) were clustered together if they showed effects in the same direction. The minimum cluster size was 3 electrodes. A cluster-level statistic was then calculated by summing the observed statistics from each sample-time point within a cluster. Next, Monte Carlo *p*-values were calculated by randomly partitioning the data 2000 times. Data were re-clustered for each permutation and a permutation distribution was constructed from the highest cluster statistics from each random permutation. Following this, we assessed statistical differences between the largest observed statistic and the permutation distribution. Resulting *p*-values were determined by the proportion of times the cluster statistics in the permutation distribution were larger than the observed statistic. The threshold for significance was set at *p* < 0.05 (one-tailed).

For comparisons that showed a significant main effect of condition, we calculated the average MI values for each participant across the electrodes and samples by which the null hypothesis was rejected in the omnibus test. Using these values, we then fit linear mixed models to assess whether the presence of syntactic *Structure* and lexical *Content* predicted the MI between EEG and annotations of syntactic node closure and node count. Models were constructed using the *lme4* package (Bates et al., 2015) in R (R Core Team, 2021), and estimates were obtained using REML and the *nloptwrap* optimizer (Baayen et al., 2008). *Structure, Content,* and their interaction were included as fixed-effects. Each factor was deviation coded, with sentence structure and English words set as reference, respectively. Random-effects structure was initially maximal, consisting of random slopes for *Structure* and *Content*, as well as by-participant random intercepts. Standardized parameters were obtained by fitting the model on a standardized version of the dataset. 95% Confidence Intervals (CIs) and *p*-values were computed using a Wald *t*-distribution approximation, with the threshold for significance set at *p* = 0.05. Potential violations of sphericity were accounted for using Greenhouse-Geisser correction. Subsequent pairwise comparisons were conducted using estimated marginal means from the *emmeans* package in R with Tukey’s method for multiple comparisons (Lenth, 2018).

### EEG Power

In addition to our evaluation of syntax tracking, we investigated whether participants evidenced differences in word-level tracking across conditions. For this, we analyzed EEG power at the rate of presentation for words (∼2Hz) because there is no word-level correlate to the structural annotations used in the analyses of MI described above due to the isochronous visual presentation. Power values were obtained using the Fourier transform on the single trial data from 0.5-2.5Hz (0.1Hz frequency resolution).

We assessed differences between conditions using a two-step process analogous to those described in the *Statistical Testing* subsection for our analysis of MI. This consisted of a cluster-based permutation omnibus test from which we chose electrodes of interest within a frequency band centered around the word-rate (1.8-2.2Hz). We then constructed mixed effects models to predict power as a function of *Structure*, *Content*, and their interaction.

## Results

In this study, we evaluated whether syntactic frames were sufficient to elicit neural synchronization in the absence of lexical content. For this, we calculated MI between abstract annotations of syntactic structure and EEG data that was recorded as participants read English sentences, jabberwocky sentences, English wordlists, and pseudoword wordlists. We additionally evaluated differences in word tracking between conditions using changes in EEG power present at the presentation rate for words (∼2Hz).

### Mutual Information for Node Closure

For our investigation of cortical tracking of node closure, cluster-based permutation testing revealed a significant main effect of condition on MI (cluster-*F* =15,496, *p* < 0.01). The cluster by which the omnibus null hypothesis was rejected consisted of electrodes with a broad scalp distribution and samples spanning the entire length of the TMIF, with peak differences spanning the lags from 0-300ms.

Following omnibus testing, we assessed whether syntactic *Structure* and lexical *Content* were statistically significant predictors of the MI values using a linear mixed model. The maximally converging model consisted of random slopes for *Content*, with by-subject random intercepts. Results indicated that the effect of *Structure* on MI was significant (β = −0.61, SE = 0.11, *p* < 0.01). No significant effects were observed for *Content* (β = 0.06, SE = 0.13, *p* = 0.67) or the interaction term (β = −0.20, SE = 0.16, *p* = 0.22). Following this, we performed post-hoc tests to evaluate pairwise differences using estimated marginal means.

Results indicated that MI for English sentences was significantly higher than that for English wordlists (Δ = 0.61, SE = 0.11, *p* < 0.01) and pseudoword wordlists (Δ = 0.75, SE = 0.13, *p* < 0.01). Jabberwocky sentences also elicited significantly greater MI than both the English (Δ = 0.66, SE = 0.13, *p* < 0.01) and pseudoword wordlist conditions (Δ = 0.80, SE = 0.11, *p* < 0.01). No significant differences were observed between English and jabberwocky sentences (Δ = −0.06, SE = 0.13, *p* = 0.67) or English and pseudoword wordlists (Δ = 0.14, SE = 0.13, *p* = 0.27). Results are depicted in *Figure 3*.

**Figure 3:**
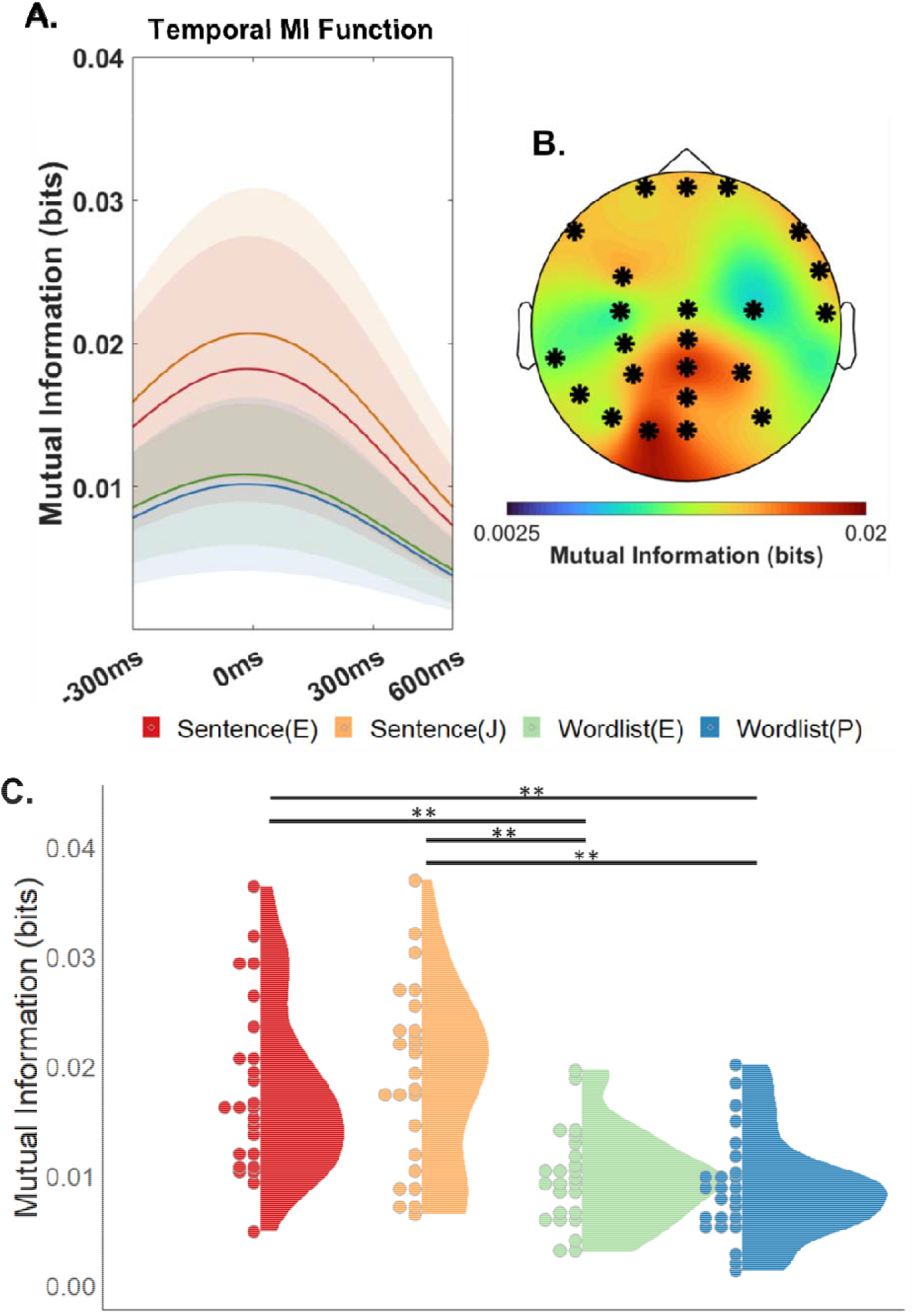
Mutual Information between EEG responses in the phrase frequency band (0.6-0.9Hz and annotations of syntactic node closure. A. Temporal Mutual Information Function. Mutual information (bits) plotted as a function of stimulus-brain lag. 0ms indicates exact alignment of syntactic annotations and EEG signal. B. Scalp topography of average MI for stimulus-brain lags ranging from 0-300ms. Highlighted electrodes comprised the cluster by which the omnibus null hypothesis was rejected during permutation testing. C. MI for English sentences, jabberwocky sentences, English wordlists, and pseudoword wordlists. Each point represents one participant’s mean MI response from 0-300ms in the electrodes identified in the omnibus test. Significant pairwise differences are denoted with ** for p < 0.05. Points were binned along the Y-axis to improve their visibility during plotting. However, all subjects evidenced distinct MI values.

### Mutual Information for Node Count

For our investigation of cortical tracking of node count, cluster-based permutation testing revealed a significant main effect of condition on MI (cluster-*F* =15,593, *p* < 0.01). The cluster by which the omnibus null hypothesis was rejected consisted of electrodes with a broad scalp distribution and samples spanning the entire length of the TMIF, with peak differences spanning the lags from 0-300ms.

We assessed whether syntactic *Structure* and lexical *Content* were statistically significant predictors of the MI values using a linear mixed model. The maximally converging model consisted of random slopes for *Content*, with by-subject random intercepts. Results indicated that the effect of *Structure* on MI was significant (β = −0.71, SE = 0.12, *p* < 0.01). No significant effects were observed for *Content* (β = 0.06, SE = 0.13, *p* = 0.66) or the interaction term (β = −0.30, SE = 0.17, *p* = 0.08). Following this, we performed post-hoc tests to evaluate pairwise differences using estimated marginal means.

Results indicated that MI for English sentences (Δ = 0.71, SE = 0.12, *p* < 0.01) was significantly higher than for English wordlists (Δ = 0.71, SE = 0.12, *p* < 0.01) and pseudoword wordlists (Δ = 0.95, SE = 0.13, *p* < 0.01). Jabberwocky sentences also elicited significantly greater MI than both the English wordlist (Δ = 0.77, SE = 0.13, *p* < 0.01) and pseudoword wordlist conditions (Δ = 1.01, SE = 0.12, *p* < 0.01). No significant differences were observed between English and jabberwocky sentences (Δ = −0.06, SE = 0.13, *p* = 0.66) or English and pseudoword wordlists (Δ = 0.24, SE = 0.13, *p* = 0.08). Results are depicted in *Figure 4*.

**Figure 4:**
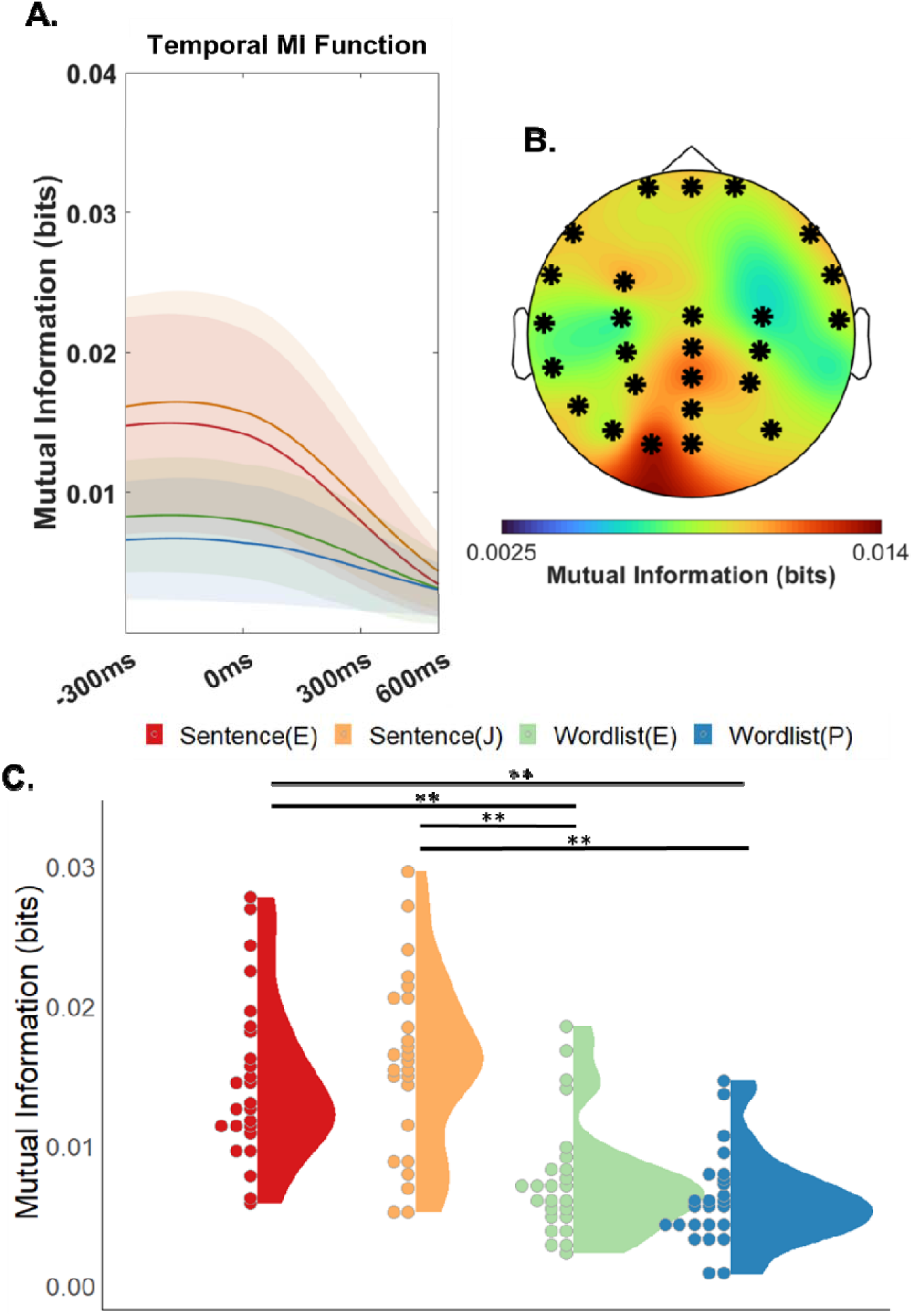
Mutual Information between EEG responses in the phrase frequency band (0.6-0.9Hz) and annotations of syntactic node count. A. Temporal Mutual Information Function. Mutual information (bits) plotted as a function of stimulus-brain lag. 0ms indicates exact alignment of syntactic annotations and EEG signal. B. Scalp topography of average MI for stimulus-brain lags ranging from 0-300ms. Highlighted electrodes comprised the cluster by which the omnibus null hypothesis was rejected during permutation testing. C. MI for English sentences, jabberwocky sentences, English wordlists, and pseudoword wordlists. Each point represents one participant’s mean MI response from 0-300ms in the electrodes identified in the omnibus test. Significant pairwise differences are denoted with ** for p < 0.05. Points were binned along the Y-axis to improve their visibility during plotting. However, all subjects evidenced distinct MI values.

### EEG Power for Word Tracking

For our investigation of cortical word tracking, cluster-based permutation testing revealed a significant main effect of condition on EEG power (cluster-*F* = 61.00, *p* < 0.01). The cluster by which the omnibus null hypothesis was rejected consisted of electrodes with a left-hemisphere scalp distribution.

We assessed whether syntactic *Structure* and lexical *Content* were statistically significant predictors of EEG power values using a linear mixed model. The maximally converging model consisted of random slopes for *Structure* and *Content*, with by-subject random intercepts. Results indicated significant effects of *Structure* (β = −0.03, SE = 0.01, *p* = 0.02) and *Content* (β = −0.04, SE = 0.01, *p* < 0.01) on EEG power. No significant effects were observed for the interaction term (β = 0.002, SE = 0.01, *p* = 0.72). We then evaluated pairwise differences using estimated marginal means.

Results indicated that EEG power for English sentences was significantly higher than that for English wordlists (Δ = 0.02, SE = 0.01, *p* = 0.02), jabberwocky sentences (Δ = 0.04, SE = 0.01, *p* < 0.01), and pseudoword wordlists (Δ = 0.06, SE = 0.01, *p* < 0.01). Jabberwocky sentences elicited significantly greater power than pseudoword wordlist conditions (Δ = 0.02, SE = 0.01, *p* = 0.04), as did English wordlists (Δ = 0.03, SE = 0.01, *p* > 0.01). No significant differences were observed between jabberwocky sentences and English wordlists (Δ = 0.01, SE = 0.01, *p* = 0.25). Results are depicted in *Figure 5*.

**Figure 5:**
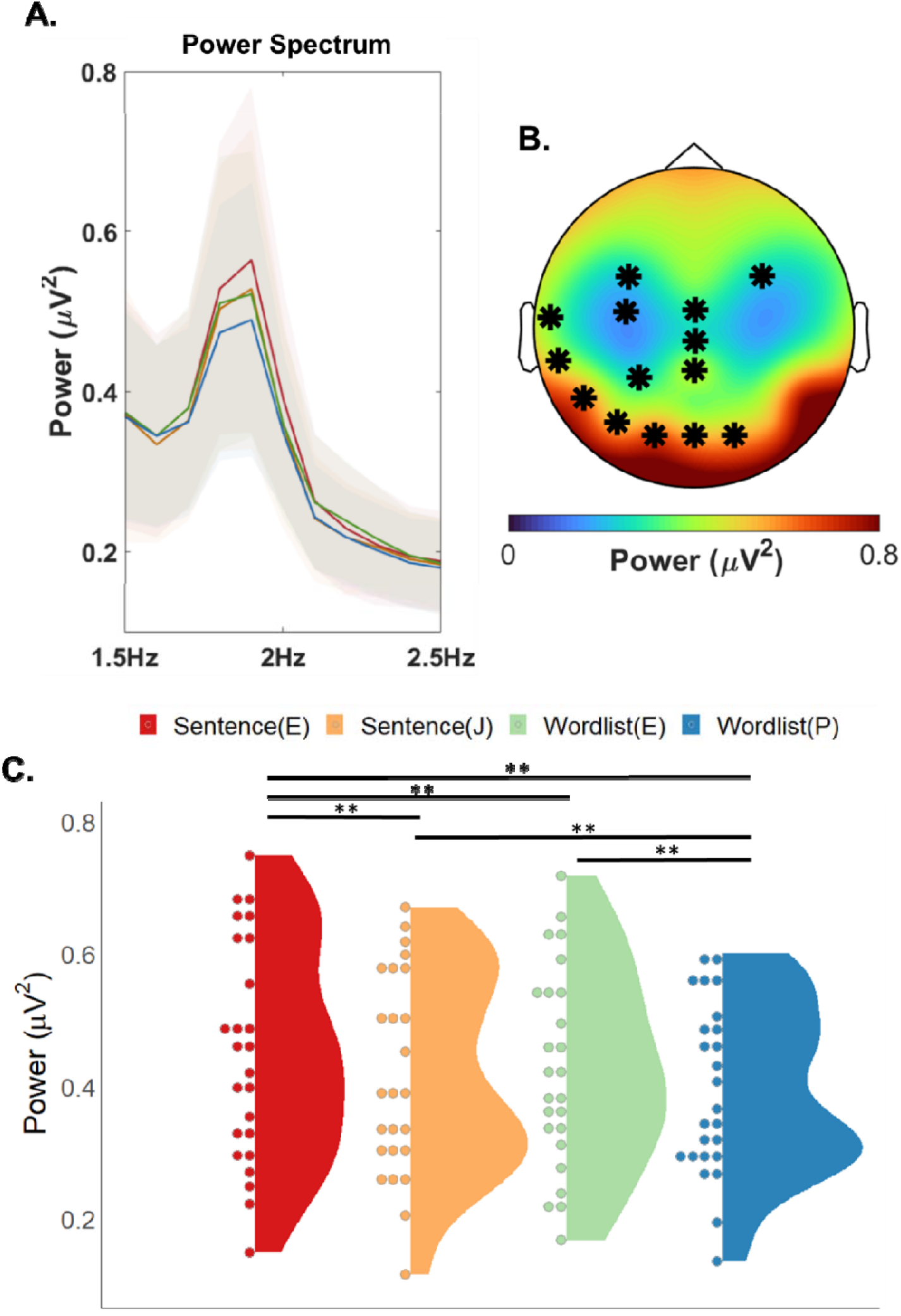
Analysis of EEG power for frequencies corresponding to the rate of presentation for words (∼2Hz). A. Power Spectrum. B. Scalp topography of power in the word frequency band (1.8-2.2Hz) averaged across conditions. Highlighted electrodes comprised the cluster by which the omnibus null hypothesis was rejected during permutation testing. C. Power for English sentences, jabberwocky sentences, English wordlists, and pseudoword wordlists. Each point represents one participant’s average power values in the electrodes identified in the omnibus test. Significant pairwise differences are denoted with ** for p < 0.05. Points were binned along the Y-axis to improve their visibility during plotting. However, all subjects evidenced distinct EEG power values.

## Discussion

In this study we set out to test whether the brain could track purely syntactic units when content words are replaced with pseudowords resulting in a linguistic pattern that would have to be inferred from the syntactic form without recourse to lexical projections. Our results are consistent with previous findings reported in Rafferty et al. (2023). Relative to wordlists that lack syntactic structure, we observed greater MI at phrase boundaries for both English and jabberwocky sentences, both of which adhere to the English caused-motion sentence structure. We found this general pattern to hold both when analyzing MI for node closures as well as when analyzing MI for node count across the sentence. MI analysis of node closures suggests that participants can track phrase-level units of English and jabberwocky, while the node count analysis suggests that participants can track the entirety of the caused-motion construction whether in English or jabberwocky.

Building on previous work by Kaufeld et al. and Coopmans et al., the results of the experiment provide further evidence that neural synchrony drives the recognition of endogenous syntactic patterns in the absence of exogenous stimuli. Our findings do diverge somewhat from these recent neuroimaging studies. In particular, Kaufeld et al. and Coopmans et al. found that jabberwocky sentences showed significantly less neural synchrony for phrases than did English sentences. Using visual presentation, in which words are easily separable from one another, seems to have eliminated this difference and enabled participants to successfully track phrases within the jabberwocky sentences. There was in fact no significant difference in neural synchrony at the phrase rate between the jabberwocky and English sentence conditions, nor at the word rate for jabberwocky and English wordlist conditions. We believe that the present results are reconcilable with the findings of Kaufeld et al. and Coopmans et al. by noting the relative difficulty of parsing pseudowords when listening to a continuous auditory stream as compared to reading sentences containing pseudowords. Indeed, Coopmans et al. acknowledge the importance of lexical parsing, although they nonetheless interpret the lack of phrase rate synchrony as evidence for a lexicalist theory of sentence comprehension. The results, therefore, suggest that phrase tracking can take place in the absence of lexically projected argument structure and would lend some support for non-lexicalist theories of sentence processing.

Several alternative interpretations of the results are nonetheless possible. First, the majority of lexicalist paradigms specify that phrasal heads project structure to syntax. Heads are generally agreed to be nouns, verbs, adjectives, and prepositions (in English at least). Some, however, have argued that NPs should better be considered as determiner phrases (DPs) (Abney, 1987). If this formulation is correct, then argument structure could have been projected from the determiners in our stimuli. While this is possible, it would not on its own explain the lack of phrase tracking in Kaufeld et al. and Coopmans et al. since jabberwocky sentences do in fact contain determiners. Moreover, the suggestion to use DPs rather than NPs does not appear to be widely accepted outside of certain generativist circles, and even Chomsky has recently indicated that the notion of DPs is unconvincing and that he believes they are “fundamentally nominal phrases” (Chomsky, 2020). Nonetheless, the present study does not rule out this possibility.

Secondly, it is possible that the difference lies in the stimulus presentation modality (i.e., reading versus listening), and this should be tested empirically. Previous research has demonstrated that phrase tracking takes place both with auditory and visual stimuli (Rafferty et al., 2023), but given that there is evidence from Kaufeld et al. that participants had difficulty parsing words from the speech stream, it seems that this is at least one important factor when considering the difference in modality. Moreover, the process of phrase tracking while listening as compared to reading must surely overlap on some level, and our findings here may be useful in helping future research isolate overlapping and diverging processes.

Coopmans et al. suggest that the parsing is accomplished “in service of semantic composition”. That is, syntax is not parsed for the sake of syntax alone, but in order to construct meaning. Similarly, several studies from the Fedorenko lab (Fedorenko et al., 2020; Mollica et al., 2020) have suggested that processing both syntax and semantics elicits a much more robust neural response in listeners. The caused motion construction used in the present study assigns clear semantic roles to the slots in the pattern which may allow participants to derive meaning from the jabberwocky sentences in spite of the lack of lexical content. It would be interesting to explore whether the same effect would be found for a syntactic pattern that did not have such obvious semantic roles.

Finally, the role of prediction should also be considered (Ferreira & Qiu, 2021; Goldberg et al., 2005). In the present study, we employed a syntactic construction whose form and meaning have high cue validity (Goldberg et al. 2005; Goldberg et al 2009), made even more predictable by being repeated multiple times over the course of the experiment, and by interleaving jabberwocky and English sentences. As a result, it is not clear whether the same results would have been found if the construction were less predictable. Indeed, it may be the case that the neural synchrony we observe is a general phenomenon that indexes the predictability of a stimulus (whether linguistic or non-linguistic). Thus, any factors of the signal that increase or decrease predictability (cue validity of the construction, cue validity of vocabulary, ambiguity etc.) may well affect measures of neural synchrony.

Taken together, the studies by Rafferty et al., the present study, and the studies by Kaufeld et al. (2020) and Coopmans et al. point to an important aspect of syntactic processing; namely, that a prerequisite of processing syntactic structures is the ability to identify the units of the structure that are to be combined (e.g., words or at the very least slots into which words conforming to the specifications of a syntactic pattern might fit). The present findings suggest that these units need not be previously known words, however. In fact, this may be an important feature of language since it allows language learners to leverage their understanding of phrase and clause level syntactic pairings between form and meaning to deduce the meaning of novel words (e.g., Fisher et al., 2010; Gleitman, 1990; Goldberg, 2006; Landau & Gleitman, 1985; Lee & Naigles, 2005). Although the present study does not negate the possibility that argument structure may project from stored lexical items, it does suggest that syntactic structures may be determined without such projections.

## Conclusion

The ability to infer syntactic representations from language input is central to comprehension. Neurophysiological evidence suggests that this ability may be supported by the synchronous firing of low-frequency neural oscillations, aligned with the occurrence of syntactic phrases. In this study, we showed that such patterns of oscillatory synchronization may be elicited by predictable syntactic templates in the absence of recognizable lexical content. These findings bridge recent neuroimaging findings with contemporary psycholinguistic data on syntactic representation.

## Acknowledgements

We thank Adele Goldberg for comments on a previous version of this paper. We also thank Cas Coopmans and an anonymous review for comments and suggestions that have greatly improved this paper. Any remaining faults in the article are, of course, our own.

## Data Availability Statement

Data are located at https://doi.org/10.5281/zenodo.8157645 and access may be provided from the authors to interested parties upon reasonable request.

