## Supplemental Table 1 for "Syntactic Constructions Drive Cortical Tracking in the Absence of Lexical Content: An Electrophysiological Investigation of Sentence Processing During Reading"

**English Sentence Stimuli**

the man pulled the boat down the dock

the hound chased the group up the path

the goat knocked the milk off the stool

the girl launched the plane through the air

the clerk hauled the box to the door

the gang kicked the stick down the road

the child rolled the truck up the street

the chief raised the gold off the ground

the snail moved the shell through the grass

the teen dropped the toad to the floor

the coach tossed the shoes down the hall

the crook lured the cop up the trail

the prince slapped the gem off the crown

the bear dragged the cub through the field

the kid pushed the bike to the school

the team flipped the rock down the hill

the guide walked the class up the creek

the champ punched the bag off the wall

the guard bumped the cart through the gate

the chef shoved the list to the side
